# *Caenorhabditis becei* recombinant inbred lines (beRILs) reveal the scope of heritable variation within a gonochoristic nematode population

**DOI:** 10.64898/2026.06.16.732751

**Authors:** Tom Parée, Jose Salome Correa, Derin Çağlar, Jacqueline L Jackson, Arielle Martel, Tuc HM Nguyen, Sophie Vallance, Matthew V Rockman

**Author notes:** Contributing authors. These authors contributed equally to this work (Tom Parée, Jose Salome Correa).

## Abstract

*Caenorhabditis* nematodes are a powerful model clade for evolutionary genetics. Isogenic lines and panels of recombinant inbred lines (RILs) are among the most essential tools for genetic studies in these species. While most *Caenorhabditis* species are gonochoristic, large RIL panels have only been developed for self-fertilizing species. This gap biases our understanding and limits our ability to address questions related to the genetic architecture of traits in outbred populations, which have radically higher genetic diversity, heterozygosity, and effective recombination than selfers. Having previously identified *Caenorhabditis becei* as a tractable gonochoristic species due to its moderate inbreeding depression, we generated two panels of advanced-intercross RILs derived from three individual outbred *C. becei* worms collected from a single locality on Barro Colorado Island, Panamá. One panel derives from a pair of worms sampled from a single rotting fig; the other derives from a cross between worms from two different figs. The panels share one founder in common, yielding two half-sib RIL panels. We sequenced and haplotyped the lines, identifying millions of variants and thousands of recombination breakpoints. Using simulations, we demonstrate the suitability of these lines for quantitative genetics studies and QTL mapping. In our single-fig panel, we observe abundant heritable variation in population growth rate, individual body size, and sexual dimorphism for body size. We detected four QTLs associated with population growth rate and show that estimated allelic effects are good predictors of selection that occurred during panel derivation.

## Introduction

Understanding how heritable variation in traits arises from underlying genotypes is a central aim of genetics. In eukaryotic organisms, natural standing genetic variation is the main source of diversity, fueling evolutionary change and providing a vast library of ecologically relevant alleles for geneticists to study. Many traits of interest in medicine, agriculture, and evolutionary biology are complex — shaped by multiple loci and environmental factors, and exhibiting continuous variation among individuals. In quantitative genetics, partitioning this variation into genetic and environmental components and mapping causal genomic regions, known as quantitative trait loci (QTL), is often the first step toward understanding molecular functions and evolutionary processes (Mackay, 2001; Stern and Orgogozo, 2008).

In many species, QTL mapping by testing correlations between alleles and phenotypic traits is performed using recombinant inbred lines (RILs). RILs are generated by (inter)crossing two or more laboratory strains or wild isolates, referred to as founders, followed by inbreeding. The resulting panel is a recombinant mosaic of founder haplotype blocks in a (mostly) homozygous state (Bailey, 1971). Homozygosity provides stable, tractable genotypes and facilitates QTL discovery by increasing additive genetic variance and exposing recessive alleles. Bi-parental panels offer a straightforward, well-controlled contrast between two parental genomes, maximizing QTL variance because alleles segregate at intermediate frequencies. As the number of founders increases, alleles are mostly present at low frequencies, but sampled allelic diversity broadens and mapping resolution improves (Pascual et al., 2016).

The self-fertile nematode *Caenorhabditis elegans* is a powerful model organism that has driven numerous advances in molecular biology, cell biology, genetics, and evolution. It grows well in laboratory culture, has a short generation time, and can be cryopreserved; its compact genome is amenable to genetic engineering, and self-fertilization facilitates the generation of inbred lines. Large collections of inbred wild isolates are now available (Crombie et al., 2024), and have been used to derive a diversity of bi-parental RIL panels (e.g., Li et al. (2006); Harvey et al. (2008); Rockman and Kruglyak (2009); Duveau and Félix (2012); Andersen et al. (2015); Greene et al. (2016)) and multi-parental RIL panels (Noble et al., 2017, 2021a; Snoek et al., 2019) that serve as important tools for QTL mapping (Evans et al., 2021; Andersen and Rockman, 2022).

Many nematode species related to *Caenorhabditis elegans* are now available in culture and have become attractive models for comparative biology, providing evolutionary context (e.g., Zauner et al. (2007); Zamanian et al. (2018); Stevens et al. (2019); Daul et al. (2019); Woodruff and Teterina (2020); Picao-Osorio et al. (2025)). While *Caenorhabditis* species vary in their mating systems and show great divergence at the DNA and protein sequence levels, they generally share morphological, ecological, and genomic features. They can be cultured under similar laboratory conditions, retaining most of the experimental advantages attributed to *C. elegans* (Nigon and Félix, 2018). However, most research has focused on only a few closely related species from the *Elegans* group, and RIL panels are available only for self-fertile species (Ross et al., 2011; Noble et al., 2021b; Stevens et al., 2022).

Deriving inbred lines in most gonochoristic (male/female) *Caenorhabditis* species has proven difficult because of extreme inbreeding depression (Rockman et al., 2025; Teterina et al., 2025). For instance, in *C. remanei*, only a few (partially) inbred lines have been generated despite substantial effort (Reynolds and Phillips, 2013; Fierst et al., 2015; Teterina et al., 2020). Deriving the CFB2252 reference strain in *C. brenneri* required approximately 100 generations of inbreeding, interspersed with regular population expansions to facilitate the purging of recessive deleterious variants (Dey et al., 2013; Teterina et al., 2025). The absence of RIL panels in gonochoristic *Caenorhabditis* species biases our understanding of genetic variation in this genus. Mating system is expected to influence nucleotide diversity and genetic architecture, including the roles of recessive deleterious variants and epistasis in shaping phenotypic variation (Wright et al., 2008; Volis et al., 2011; Chelo et al., 2014, 2019; Cutter, 2019; Clo et al., 2021; Abu-Awad and Waller, 2023). The lack of gonochoristic RIL panels also limits opportunities for studying outcrossing-related traits that are absent or only vestigial in predominantly selfing species.

We previously identified *C. becei*, a species of the *Japonica* group, as a tractable gonochoristic *Caenorhabditis* species due to its moderate inbreeding load, with a relatively large number of lines surviving 20 successive generations of sib-mating (Rockman et al., 2025). We have begun developing tools to study *C. becei*, including the isolation of wild populations from Barro Colorado Island, Panamá (Sloat et al., 2022), and the generation of inbred and transgenic lines and a chromosome-level genome assembly (Salome Correa et al., 2025). *C. becei* retains all of the laboratory conveniences of *C. elegans*, including culture conditions and cryopreservation. They differ in mating system, optimal growth temperature (25°C rather than 20°C), and developmental rate, which proceeds egg to egg in less than 40 hours under our standard conditions. As we show below, *C. becei* also differs significantly from *C. elegans* in its levels of genetic diversity.

In the present study, we report the construction and analysis of two RIL panels. Each RIL panel originates from two non-inbred wild individuals and is therefore composed of four founding haploid genomes. The two panels share one common founder and are thus derived from two half-sibling families. This design was specifically chosen to facilitate studies of inbreeding depression by regenerating F1 outbred genotypes from crosses between RILs harboring varying degrees of genetic similarity. Here, we introduce these panels as tools for studying quantitative genetics in a gonochoristic *Caenorhabditis* species, using both simulated and experimentally generated data. We show that a small sample of *C. becei* diversity from a single locality – indeed, from a single rotting fig – contains enormous stores of sequence variation that tags genetic contributors to heritable variation in body size, sexual dimorphism, and population growth rate. The latter is a fitness proxy, and QTLs for it explain subtle allele frequency shifts that occurred during strain construction.

## Materials and Methods

### Culture conditions

If not stated otherwise, nematodes were cultured at 25°C on 10cm nematode growth medium (NGMA) plates (1% agar and 0.7% agarose), with *E. coli* (*OP50-1*) as a food source. Whenever needed, nematode samples could be synchronized by bleaching or cryopreserved following Stiernagle (2006).

### Founder sampling & RIL panels derivation

Source populations were isolated from fallen figs collected along the Lathrop Trail, Barro Colorado Island, Panamá (9.164°,-79.844°), on August 25 (QG2990) and August 26 (QG2987), 2018 (Sloat et al., 2022). Worms were extracted from the samples by placing the fig directly on an NGM plate (QG2987) or by Baermann funnel onto seeded NGM plates (QG2990) (Tintori et al., 2022). Sample QG2990 contained tens of thousands of *Caenorhabditis* nematodes, including adults, larvae, and dauers. Sample QG2987 contained hundreds of *Caenorhabditis* adults and larvae. The entire populations were returned to New York, where they were passaged by chunking, cleaned of bacteria by bleaching, and then cryopreserved on September 7, 2018. Thus, in each case, the populations experienced several generations at large population sizes between collection and cryopreservation.

A male and a female, referred to as founder *FM* and founder *Fα* were picked from thawed samples from the QG2990 population. A second female, referred to as founder F*β*, was picked from thawed samples from the QG2987 population. To initiate two distinct panels, the male founder *FM*, was successively mated with the two isolated females (Figure 1). To allow for recombination of founder haplotypes, each cross was expanded separately for 9 days with regular chunking to avoid starvation, corresponding to approximately 4 to 5 generations before being cryopreserved. To generate RILs, samples from each cross were thawed, and from crosses *α* and *β*, respectively, 351 and 323 mating pairs were picked and passaged by sib-mating for 25 generations. In the first 10 generations, failed crosses were abandoned; in later generations, we sometimes rescued failed crosses by retrieving animals from the previous generation’s plate. 189 of the 674 lines survived 25 continuous generations of inbreeding (100 *α* and 89 *β*), and an additional 108 lines (58 and 50) survived 25 generations after rescues, yielding a total of 297 RILs generated. Table S1 contains a summary and description of the lines and populations.

**Fig. 1.**
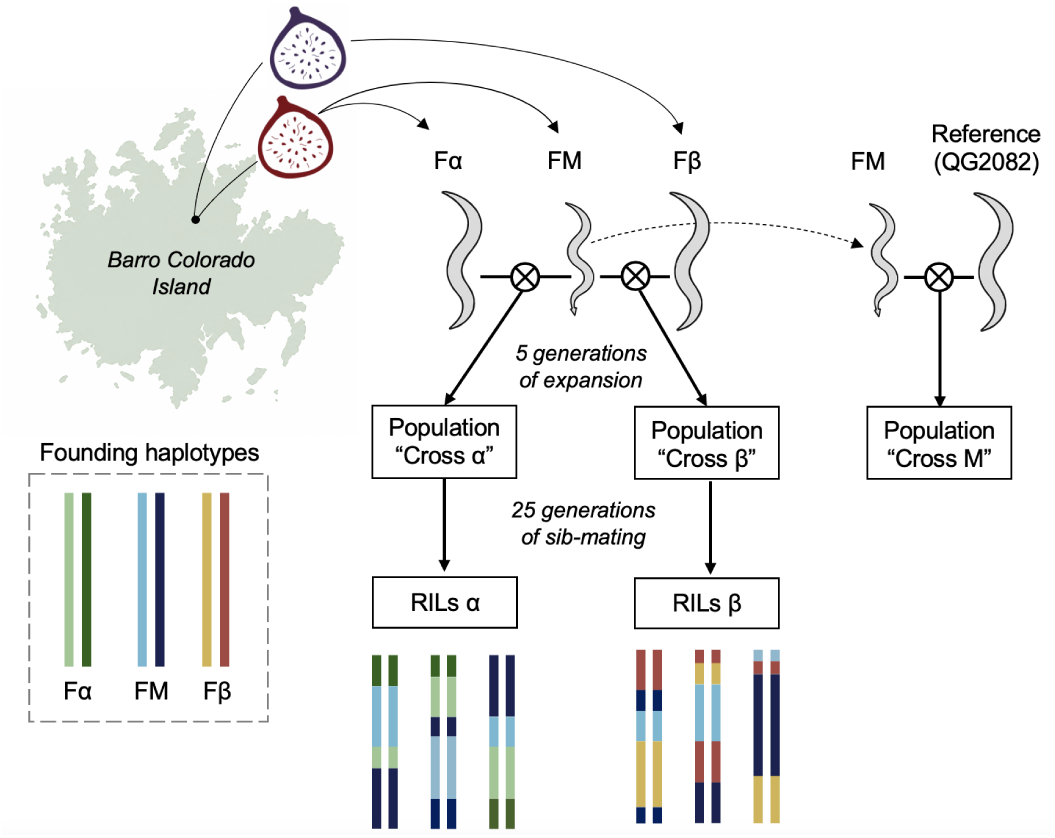
Panel derivation design. Three founders were sampled from fallen figs at a single location on Barro Colorado Island. F*α* and FM were derived from the same fig, whereas founder F*β* was derived from a different fig. Founder FM was male and was mated with the two other founders, which were females. These crosses produced two half-sibling families that were expanded for ∼5 generations. Recombinant inbred lines were subsequently derived through sib-mating. Founder FM was also mated with a female from the QG2082 reference strain solely to facilitate phasing of the founder haplotypes (see Methods).

To facilitate the subsequent reconstruction and phasing of the founder haplotypes, the founder M was crossed with the QG2082 reference strain. Expanded populations, denoted “Cross M”, were pool-sequenced. Samples of population *α* and *β* were pool-sequenced before cryopreservation and after cryopreservation at higher depth.

### Library preparation & sequencing

DNA libraries were prepared from total DNA extraction and sequenced for the expanded populations and the RILs. DNA libraries were prepared using the NEBNext Ultra II FS DNA Library Prep Kit for Illumina (New England Biolabs) with an input of 200ng of DNA per sample; quantified by Qubit dsDNA HS Assay Kit (Thermo Fisher Scientific). Fragmentation was performed by incubation at 37°C for 10 minutes, followed by 65°C for 30 minutes, with the thermocycler lid set to 75°C. Indexing was performed using NEBNext Multiplex Oligos for Illumina (96 dual indices per plate, 4 plates total). Ligation was carried out at 20°C for 15 minutes. PCR enrichment consisted of an initial denaturation at 98°C for 30 seconds, followed by 5 cycles of 98°C for 10 seconds and 65°C for 1 minute and 15 seconds, with a final extension at 65°C for 5 minutes. Size selection and purification of the libraries were performed using NEBNext Sample Purification Beads, selecting for fragment sizes of 400–600bp. Libraries were eluted in DNA Elution Buffer (10mM Tris-HCl, pH 8.5, 0.1mM EDTA), quantified using the Qubit dsDNA HS Assay Kit, and a subset was analyzed for fragment size using the Agilent TapeStation D1000. Libraries were normalized and pooled at equimolar concentrations. Final pool quality was confirmed using TapeStation D1000 and quantified using the KAPA Library Quantification Kit for Illumina platforms. Libraries had an average insert size of ∼500 bp.

Libraries were sequenced across multiple sequencing runs between 2019 and 2025 at the New York University GenCore on an Illumina NovaSeq 6000 platform (S2 or S4 flow cells; 2 × 150 bp paired-end; 300-cycle v1.5 chemistry), except for the population samples collected prior to cryopreservation, which were sequenced on an Illumina NextSeq 500 platform (1 × 75 bp single-end; 75-cycle v2.5 chemistry). The realized sequencing depth was 17× and 218× for the expanded population samples prior to and after cryopreservation. The realized median depth across the RILs was 41×, and the average depth is higher than 5 in 93% of RILs. The GenCore staff used Picard IlluminaBasecallsToFastq (version 2.23.8; https://broadinstitute.github.io/picard/) for base calling, with APPLY_EAMSS_FILTER set to false, and Pheniqs version 2.1.0 for demultiplexing (Galanti et al., 2021).

Reads were aligned to the *Caenorhabditis becei* reference genome (strain *QG2082*, assembled in Salome Correa et al. (2025)) using bwa mem (Li and Durbin, 2009). Aligned reads were filtered using samtools (Li et al., 2009) to retain properly mapped alignments with mapping quality ≥20 (-q 20), while excluding unmapped reads (-F 0x4), reads with unmapped mates (-F 0x8), and secondary alignments (-F 0x100). Variant calling was performed using bcftools mpileup (Danecek et al. (2021); -q 30 -Q 20) and choosing genotype that minimises the Phred-scaled genotype likelihoods (PL), yielding a total of 5,078,688 variants with QUAL ≥ 20. A very stringently filtered dataset was also created as a base to reconstruct the founding haploid genomes using only the 521,138 most confident diallelic SNVs (QUAL=999 & MQ ≥ 59 & AC*>*1 &MQB*>*0.001 & BQB *>* 0.001 & MQSB *>* 0.001 & DP*>*20000 & DP*<*30000 & missing *<* 10 & het *<* 15).

### Variant filtering and haplotype calling

Given that the RILs are a recombinant mosaic of six founding haploid genomes, the best strategy to filter variants is arguably to call haplotypes and keep compatible variants. Although the six founding haplotypes are unobserved, they can be confidently reconstructed by leveraging the allele frequencies in the three populations (*α, β, M*) and the linkage present in the RILs. To reconstruct haplotypes, we developed a custom method tailored to the specific features of our dataset. The method is described in detail in Appendix. In total, six founding genomes were phased for 2,442,980 variants, including 276,288 that are fixed for the non-reference allele (all six founder haplotypes differ from the reference genome at these sites). The excluded unphased variants have lower quality control metrics and a more variable depth distribution, indicating that these generally are lower quality calls. Of the remaining 2,166,692 phased variants segregating in our crosses, 276,288 are fixed alternative alleles, 50,300 variants are multi-allelic, 347,614 likely overlap a deletion (always missing in one founding haplotype), and 211,481 likely tag a duplication (always heterozygous within a reconstructed haploid founding genome and depth distribution skewed towards higher values). After filtering out these categories, 1,557,297 variants are retained for downstream analysis.

### Simulation of panel derivation

Individual-based simulations of the panel derivations were performed using SLiM 4.0.1 (Haller and Messer, 2023), using a non-Wright-Fisher model for greater flexibility. Simulations were run separately for each panel and each chromosome. Each simulation began with a cross between two founders. To track haplotype identities, 500 neutral markers, evenly spaced by genetic distance, were used to tag each haploid genome, with a unique allele assigned to each one. Because there are four founding haploid genomes but SLiM only supports biallelic loci, each marker was implemented as two perfectly linked biallelic loci, allowing the representation of four distinct allelic combinations. Following the initial cross, the population underwent an outcrossing phase. Although the experimental outcrossing phase involved population expansion and overlapping generations with individuals of different ages, this complexity is not well characterized and was therefore simplified in the simulation. Specifically, the outcrossing phase was modeled as a few non-overlapping generations at a constant population size. We varied the population size (*N* = 400 or 4000) and the number of generations (4 to 7) to examine how uncertainty in the outcrossing phase parameters might influence the resulting RIL panels. After the outcrossing phase, mating pairs were randomly selected and isolated to initiate 25 generations of sib-mating, mimicking the derivation of inbred lines. Simulated RIL haplotype information was extracted and analyzed in R (R Core Team, 2024). The haplotype tag dataset was converted into an SNV matrix by assigning experimental SNVs according to the founder identity of each haplotype block. This approach allowed us to mimic the experimental data and account for similarities among founding haploid genomes that may have rendered some crossovers effectively invisible. Finally, for each RIL, we assessed the number and positions of recombination events and heterozygous diplotypes. For each parameter set, 50 simulation runs were performed.

### Phenotyping

Resource exhaustion time was used as a proxy for multi-generational population growth rate using the scanner assay developed by Tintori et al. (2025). Specifically, we measured the number of hours before starvation in an expanding population, a metric that increases as the growth rate decreases. For each strain, two virgin adult females and two males were placed on 3.5 cm petri dishes containing NGM agar and a lawn of *E. coli* previously killed by paraformaldehyde. The plates were incubated at 25°C and imaged every hour, which allows detection of the time point at which the population peaks before starvation. A total of 154 strains were phenotyped across nine experimental blocks, with an average replication of 3.2 per strain and 27 strains phenotyped in two blocks. The distribution of the number of hours before starvation across strains is exponential and has an inverse relationship with growth rate; therefore, its negative logarithm was used for downstream analyses.

Sex-specific size was defined as the average individual length 48 hours after the L1 stage. Within each experimental block, strains were bleached simultaneously, and the resulting eggs were left to hatch overnight in M9 buffer at 240 rpm and 25 °C. L1 larvae were then seeded onto fresh petri dishes and allowed to develop at 25°C for 48 hours, after which the plates were washed with 10 mL of M9. Samples were centrifuged at 4 × g for 2 minutes, and the supernatant, containing L1 larvae and unhatched eggs, was discarded, retaining only the pellet, which primarily consisted of adults. A subset of each sample was dispensed into 12 wells of a 96-well plate (Fisher Brand, FB012931), at an average density of fewer than 50 adults per well, in a volume of 65 *µ*L. To immobilize the worms for imaging, 330 *µ*L of 50 mM sodium azide was added to each well, and plates were incubated at room temperature for at least 10 minutes. Imaging was performed in brightfield mode with a 20 ms exposure using the Molecular Devices ImageXpress Nano microscope (Molecular Devices, San Jose, CA) with 2X magnification. Individual traits, including body length and curvature, were extracted following Widmayer et al. (2022), processing the images through the CellProfiler’s WormToolbox (Wählby et al., 2012) and the easyXpress R package (Nyaanga et al., 2021). Objects were classified by sex using an eXtreme Gradient Boosting (XGBoost) model [xgboost R package, Chen et al. (2015)]. We first trained the model on 23 features extracted from 3,532 manually classified objects (female: n=2,648; male: n=884). Model performance, evaluated by training on 90% of the data and testing on the remaining 10%, showed 94% correct sex classification, 2.5% incorrect, and 3.5% unclassified. In total, 144 strains have been phenotyped across 7 blocks. For each replicate, mean and standard error for each sex is calculated and used for downstream analysis.

For plotting Figure 4A,B, we estimated marginal means of strains’ phenotype for each trait, obtained from a linear mixed model [lme4::lmer and emmeans::emmeans R functions; Lenth (2024); Bates et al. (2015)] including a random block effect, and, for size only, a fixed effect for sex:

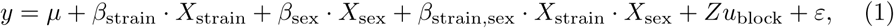

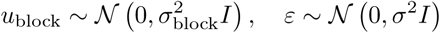

where *y* is the vector of phenotypes, *µ* is the intercept, and *β* are the fixed-effect coefficient for the variables *X*. The term *u*_block_ is the random block effect. The third and fourth terms containing the *X*_sex_ variable are only modeled for size.

### Sexually convergent and sexually divergent size components

Male and female size can be treated as two distinct yet correlated phenotypes measured in the same strains. In multi-trait analyses, canonical transformation and related methods are commonly used to reduce the dimensionality of mixed-model equations (Ducrocq and Besbes, 1993). The goal of such transformations is to convert n correlated traits into n uncorrelated linear combinations, allowing independent statistical testing without loss of information. These transformed traits define orthogonal axes of the phenotypic space. In the case of two mean-centered and scaled traits plotted on Cartesian coordinates, the orthogonal phenotypes can be obtained by rotating the data by 45°, such that the two transformed trait values respectively correspond to the x and y coordinates of the rotated system. For sex-specific traits, this transformation is particularly meaningful: the two rotated traits represent the sexually convergent and sexually divergent components of variation (e.g., Ruzicka et al. (2019)).

### SNV-based heritability

Genomic heritability 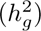 of a trait in a given population can be estimated by decomposing the phenotypic variance 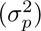 into the variance explained by genetic variation captured by genome-wide molecular markers 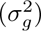 and the residual variance 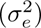 (de Los Campos et al., 2015):

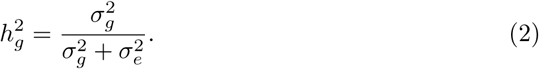

Estimating these variance components can be achieved by measuring phenotypes in a genotyped reference population (here, the RILs *α*) and fitting linear mixed models (LMMs) that incorporate genomic correlations between individuals through a genomic relationship matrix (GRM):

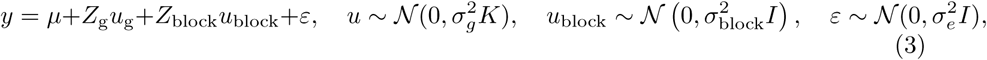

where *y* is the vector of phenotypic observations, *µ* is the intercept, *Z* are the design matrices for the random genetic effects *u_g_*and the random block effect *u*_block_, and *K* is the GRM of dimension *n* × *n*, with *n* the number of strains. *K* can be obtained from the genotype matrix *M*, of dimensions *n* × *l*, where *l* is the number of loci. Using the widely adopted method of VanRaden (2008) to estimate *K*, the matrix is centered around 0, and the average of the diagonal elements, representing self-relatedness, is expected to be 1 under Hardy-Weinberg equilibrium, but 2 for inbred lines (i.e., 1 + *f*, where *f* is the inbreeding coefficient (Endelman and Jannink, 2012)). It is generally recommended to use the standardized *K* with an average diagonal of 1 (Speed and Balding, 2015; Feldmann et al., 2022). This recommendation is not applicable when the reference population is used to estimate genetic variance in another related population with a different genetic structure (Legarra, 2016). This is not our case, as we aim to directly estimate 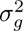 observed in the reference RILs population, the quantity relevant for QTL mapping. We thus use the average semi-variance (ASV) method to directly compute a standardized *K*:

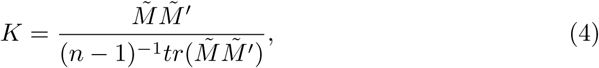

where *M̄* is the mean-centered marker matrix, and tr() is the trace of the matrix (i.e., the sum of diagonal elements). Estimates of heritability were implemented in R using the sommer::mmes function to fit the LMM (Covarrubias-Pazaran, 2016). For genomic variance partitioning (Yang et al., 2011), a GRM was separately computed for different genomic intervals and fitted into the same LMM. As linkage disequilibrium (LD) can bias genomic variance estimates because linked markers do not provide independent information about the genetic relationships (Speed et al., 2012), the genotype matrix *M* was pruned to exclude fully redundant markers (*r*^2^ *>* 0.999), resulting in a dataset containing 116,217 SNVs.

### Haplotype-based heritability

Heritability was also calculated from the variance components estimated using relatedness derived from haplotype information. Following Hickey et al. (2013), we computed the haplotype-based genomic relationship matrix, *G_HAP_* _1_ (following their notation), which accounts for varying similarity between haplotypes. The genome was divided into 1 cM windows, and *G_HAP_* _1*,i*_ was independently calculated for each *i*^th^ window. The *h* different haplotypes are listed, and a *h* × *h* haplotype similarity matrix, denoted *H*, is calculated. The element *h*1*, h*2 of this matrix correspond to the haplotype similarity score, defined as the proportion of SNVs that are identical between the *h*1 and *h*2 haplotypes. The relationship between the *j*^th^ and *k*^th^ strain at the *i*^th^ window is then calculated as:

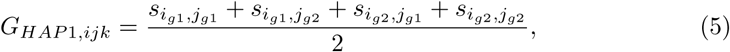

where *s_i__g_*_1_*,j_g_*_2_is the haplotype similarity score between the first haplotype of the *i* strain and the second haplotype of the *j*^th^ strain. *G_HAP_* _1_ is then obtained by averaging all the *G_HAP_* _1*,i*_. This results in a relationship matrix that is not standardized, where self-relatedness of an inbred individual is 2 and genetic relationship values are always positive. The relationship values are closer to a coancestry coefficient and should be regarded as relationships relative to an unobserved population of unrelated individuals. The resulting estimate of genomic variance is therefore not directly comparable with that obtained from SNV-based standardized genetic similarity relative to the reference population. Legarra (2016) presents a simple method for rescaling 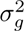 obtained using any relationship matrix, standardized or not, into the genomic variance of the reference population, 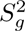:

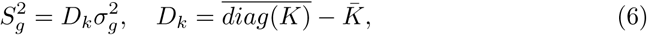

Where *K̄* is the mean of *K̄* and 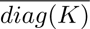 is the mean of its diagonal elements. To then obtain haplotype-based heritability presented in the main text, 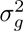 was replaced by 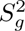 in Equation 2.

### Single marker mapping

For a given trait, single-marker QTL mapping was performed using the EMMAX algorithm (Kang et al., 2010) that we implemented in R with the leave-one-chromosome-out (LOCO) approach (Yang et al., 2014):

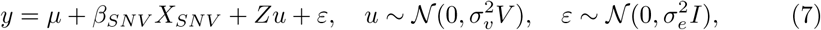

where *X_SNV_* is the matrix of SNV genotypes and *β_g_* is the effect of SNVs on phenotypic variation that is estimated. V is the covariance matrix, which is the kinship matrix, *K*, when there is no other random effect. To remain consistent with previous equations where we treated block as a random effect, we obtained a composite covariance matrix by combining the kinship matrix and the block covariance matrix (1 when replicate i and j are assayed in the same block, 0 otherwise), weighted proportionally to their respective variance components estimated in equation (3). However, note that using *V* = *K* and treating block as a fixed effect does not change our conclusions about the discovered QTLs. Because we use the LOCO approach, *K* is recalculated for each chromosome, excluding the chromosome of interest.

To determine significance thresholds, we employed a permutation-based approach (John et al., 2024). Specifically, we randomly permuted the genotypes across strains at least 500 times, conducting a QTL mapping for each permutation. Genotypes are permuted after kinship matrix calculation to not lose population structure and the relationship between phenotypic observations. For each iteration, we recorded the smallest *p*-value obtained. The significance threshold corresponding to a desired *α* error level was then established as the *α* quantile of the distribution of these minimal *p*-values across all permutations.

To enhance computational efficiency, mapping on unpermuted and permuted dataset were performed using genotype data with redundant markers (*r*^2^ *>* 0.999) excluded. As these markers are fully redundant, no information was lost in this process.

### Haplotype QTL mapping

For a given trait, a haplotype QTL mapping was performed by iterating across 1 cM windows, and resolving the following LMM [sommer::mmes() R function Covarrubias-Pazaran (2016)]:

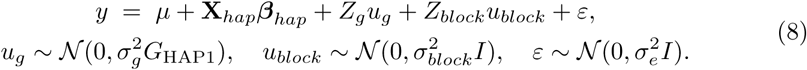

With **X***_hap_* being the indicator matrix used to encode the haplotypes block identity as a multi-level categorical variable, and other variables defined as above. Haplotype levels were defined as the founders’ haploid genome identity at the mid position of the window. If some founders’ haploid genomes are identical within a given window, they are merged into a unique level.

### Power to detect a QTL

The power to detect a QTL was estimated by implementing simulations where the additive genetic variance 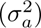 is half of the phenotypic variance 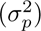, so *h*^2^ = 0.5.

Causal loci were sampled from the pruned SNV dataset. One random SNV was defined as the QTL of interest, with its allelic effect size set such that the variance explained by the QTL 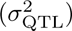 was equal to a specified proportion of the total phenotypic variance. Additionally, 1,000 other causal loci represented a polygenic background, with allelic effects drawn from a normal distribution with variance equal to the remaining additive genetic variance 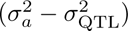. QTL mapping was performed as described for the equation (7), without the sex-specific term. The significance threshold for QTL detection was determined by performing QTL mapping on 1,000 simulated datasets in which strain phenotypes were randomly sampled from a normal distribution with variance 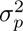, and thus containing no genetic effects 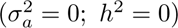.

### Data availability

Genotype tables, phenotypic data, and scripts used to conduct the analysis are available at the associated Github repository, and will be deposited on Dryad upon publication. Sequencing data will be shared on NCBI upon publication of this article. RILs and populations are available on request.

## Results

### Founder collection and panel derivation

We set up matings between a single male (FM) and two females (F*α* and F*β*) derived from wild-caught populations isolated from fallen figs, resulting in two full-sibling families sharing the same father (Figure 1). Founders F*α* and FM originated from the same fig, whereas founder F*β* originated from a population isolated from another fig located ∼30 m away. Each family was then expanded to a population size of several hundred thousand individuals over approximately five generations, after which the populations were cryopreserved and sequenced. Starting from 676 random mating pairs across the two expanded families, we generated 297 *C. becei* recombinant inbred lines (beRILs) through 25 successive generations of sib-mating. Among them, 189 lines were never rescued at any point, meaning that sib-mating was continuously successful throughout the derivation process (see Methods). RILs were successfully genotyped at 2.2 million high-quality variants segregating genome-wide, of which 1.55 million informative di-allelic markers were retained for downstream analyses. Excluded variants included, for example, multi-allelic SNVs or variants likely overlapping a deletion and missing in one or more founding haplotype (see Methods for details).

### Nucleotide diversity, population structure and linkage disequilibrium

With six founding haplotypes in total, most SNVs segregate at intermediate frequencies among the beRILs (Figure S1A). No SNVs were lost during RIL derivation, yielding a density of 17 SNVs/kb. As expected from the “two half-sib panels” design, PCA highlights substantial population structure among all RILs (Figure S1B), with panel *α* containing 2.7 SNVs/kb and panel *β* 4.8 SNVs/kb that are fixed in the other panel. SNV density is slightly lower in RIL panel *α* compared to panel *β*, with an average of 12 and 14 SNVs/kb, respectively. Despite the population structure arising when pooling the two panels, intra-chromosomal linkage disequilibrium (LD) is lower across panels than within each panel, reaching *r*^2^ = 0.3 between alleles separated by 0.5 cM (Figure S1C,D). The overall reduction in LD observed when pooling panels is attributable to the increased haplotype diversity. As expected, LD is substantially higher in the lowly-recombining center domains than in the highly-recombining arm domains (Figure S2).

We did not detect excess in the mean inter-chromosomal LD for any chromosome pair (Figure S3). This contrasts with panels derived from geographically distant *C. elegans* strains, where such excess LD has been observed, likely reflecting epistatic interactions (CeMEE; Noble et al. (2017)). The emergence of long-range epistatic interactions, including genetic incompatibilities and co-adapted loci, is expected to be favored by self-fertilization and reproductive isolation (Colomé-Tatché and Johannes, 2016; Nguyen et al., 2025). The absence of such long-range LD is thus consistent with the genetic architecture of an outbred local population.

### Founding haplotypes and recombination

Founders’ phased haplotypes were reconstructed from alleles and haplotypes segregating in the expanded populations and RILs. Founders F*α* and FM share large identical by descent (IBD) regions, particularly on chromosomes I, IV, and X (Figure S4). They also exhibit longer runs of homozygosity than founder F*β*, resulting in a higher estimated inbreeding coefficient (*f_ROH_*) (Table S2). These explain the slightly reduced SNV density in the panel *α*, mentioned above, and are consistent with the fact that founders of the panel *α* were isolated from the same fallen fig. Elevated homozygosity within a fig is consistent with colonization of a new food patch by a restricted number of individuals, leading to inbreeding and population structure. Moderate inbreeding in *C. becei* may have favored the purging of strong-effect recessive deleterious alleles, which could explain why no founding haplotype was lost during inbreeding in the beRILs and why *C. becei* suffers only mild inbreeding depression (Rockman et al., 2025).

Founding haplotype identities were inferred across the genome of RILs (Figure 2A). All founder haplotypes are represented across the two panels, with the minimum founder haplotype frequency within a panel across all genomic positions being 7% (Figure 2B). The average change in frequency from the founders during panel construction is only 5.6 %. This suggests that highly-penetrant medea toxin–antitoxin elements, known to segregate among other unrelated strains (Salome Correa et al., 2025), and which can cause strong segregation distortion, are not polymorphic in this panel.

**Fig. 2.**
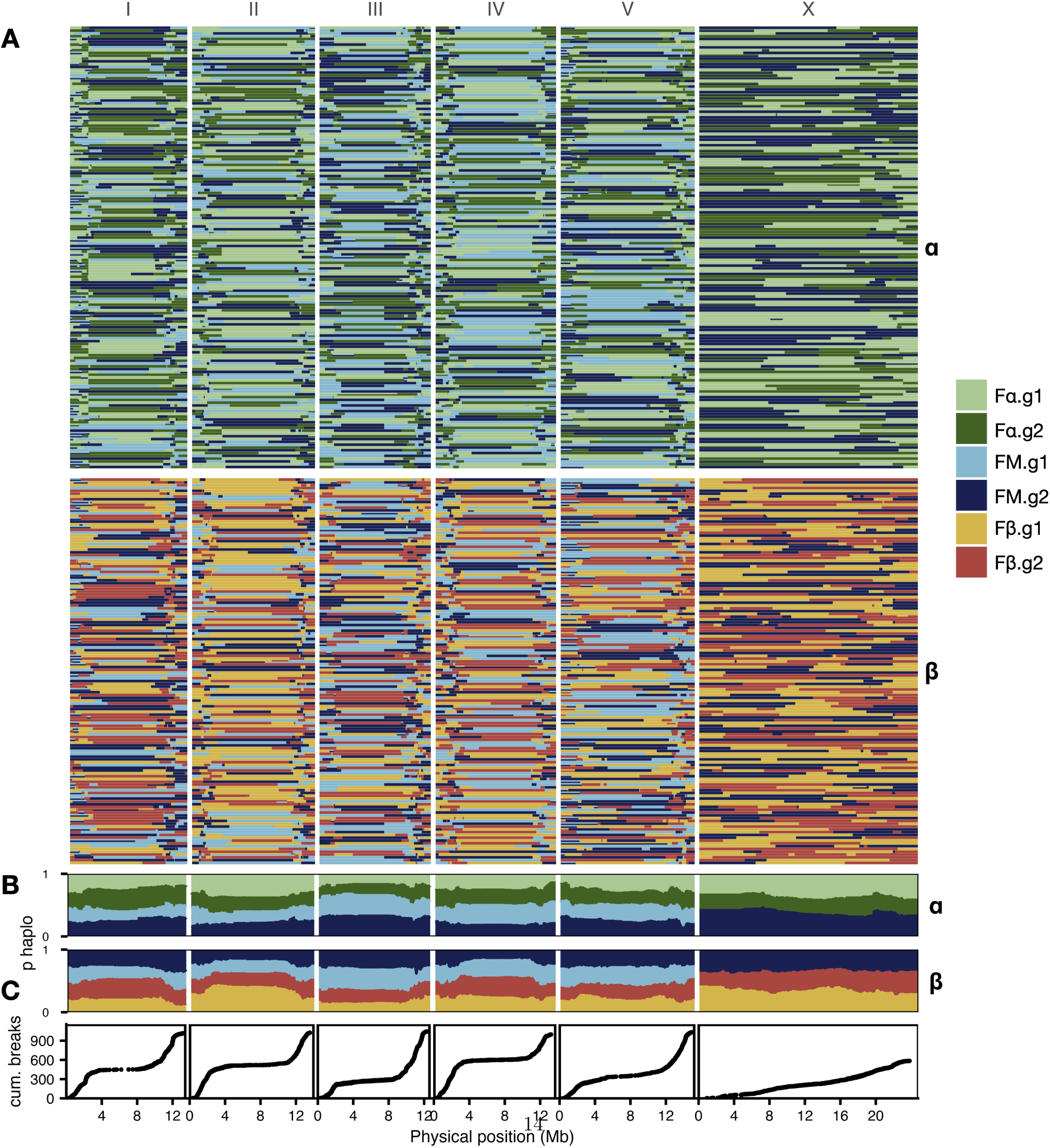
RIL haplotypes. **A.** Each row represents one RIL, and colors indicate the founder origin of genomic segments. **B.** Proportion of the founding haplotype across the genome. Pairs or light/dark colors indicate founding haplotypes originating from the same founder (green: *Fα*, orange: *Fβ*, blue: *FM*; same color coding as Figure 1). **C.** Cumulative distribution of recombination breaks across the chromosomes.)

Segments from all four haploid genomes founding a given panel are present in every RIL. A given RIL’s haploid genome is composed of an average of 24.3 founding haplotype blocks for the panel *α* and 27.5 for the panel *β*. This difference is expected given the longer IBD regions between founding haplotypes, leading to a higher proportion of uneffective recombination events in panel *α*. Recombination breakpoints are unevenly distributed along chromosomes, being rare in the center (Figure 2C), as expected from the recombination landscape in *C. becei* and other *Caenorhabditis* species (Salome Correa et al., 2025). In total, 5,696 recombination breakpoints were detected, with an average physical distance between consecutive breakpoints of 8.2 kb in chromosome arms and 46.0 kb in chromosome centers.

The breakpoint distribution recovers broad-scale characteristics of the *C. becei* recombination landscape, including the exceptionally high recombination rate on the right arm of chromosome III (Salome Correa et al., 2025). Neutral individual-based simulations show that the relative distribution of recombination breakpoint positions closely matches expectations given the previously published genetic map (Figure S5A). An exception is the left arm of chromosome I, where an excess of crossovers is observed in panel *α*, whereas fewer crossovers than expected are observed in panel *β*. Such patterns could result from selection for or against specific recombinants in this genomic interval. Alternatively, they could reflect segregation of distinct recombination modifier alleles between the two panels.

A slightly higher total number of recombination breakpoints is observed in the neutral simulations than in the experimental data (Figure S5B). This difference may simply reflect an artifact stemming from simulating non-overlapping generations, whereas generations overlapped during panel derivation, allowing less-recombined individuals from earlier generations to continue contributing to population expansion at the end of the outcrossing phase.

### Residual heterozygosity

Despite 25 generations of inbreeding, the RILs have maintained, on average, 0.45% of heterozygous SNVs. Heterozygous diplotype blocks are detected in 59 RILs in panel *α* and 53 RILs in panel *β* (Figure S6). The number of heterozygous RILs does not differ from neutral simulation, showing this level of heterozygosity is expected given the number of generations of sib-mating (Figure S5C). When present, the typical heterozygous region is small, spanning on average 1.8% of the physical genome size in these lines. No region is overrepresented among the heterozygous segments, and every founder’s haplotype is represented among the RILs in a homozygous state across all genomic positions. Therefore, our results suggest that strongly (pseudo-)overdominant loci or recessive lethal alleles did not segregate in the founders.

### Power to detect a QTL

We estimated the power to detect a single QTL in a polygenic background using simulated datasets with 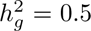. When using all RILs, the detection power at error rate *α* = 0.05 reaches 70% for QTLs explaining as little as 12% of the phenotypic variance (Figure 3A). At this level of variance explained, the causal variant is on average 0.5 cM from the position with the lowest *p*-value (Figure 3B). Power and mapping resolution decrease when using a single panel; for panel *α*, for example, 70% detection power is only reached when the QTL explains 18% of the variance (Figure S7B).

**Fig. 3.**
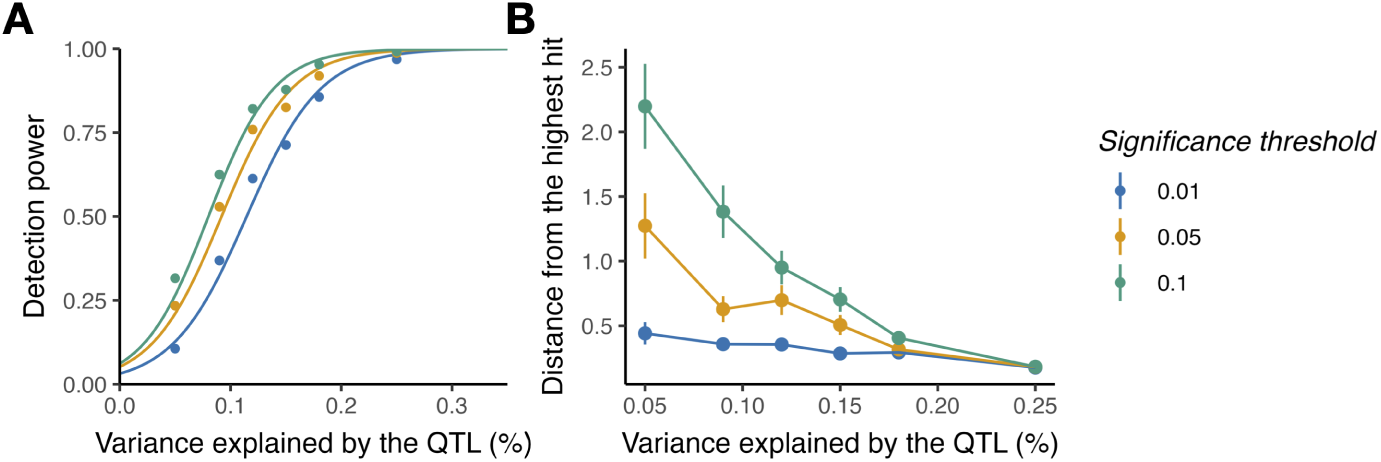
Power analysis. Phenotypic data for a trait with *h*^2^ = 0.5 were simulated across RILs by randomly sampling a single major QTL and a polygenic background from the experimental SNVs. For varying proportions of phenotypic variance explained by the QTL, we assessed (**A**) the proportion of simulation runs in which the QTL was detected and (**B**) the distance between the detected peak and the true QTL.

### Heritability of growth rate and sex-specific size

We experimentally validated the suitability of the beRILs for quantitative genetics by phenotyping panel *α* RILs for two traits. The first trait is the multi-generational population growth rate, measured using food consumption rate as a proxy and expected to be strongly influenced by brood size and egg-to-adult developmental time (Figure 4A). Large differences were observed between strains, with the fastest growing strain reaching the pre-starvation time-point in 3.4 days, whereas the slowest required 9.4 days. We find high genomic heritability 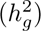 for multi-generational growth rate, when estimated using genomic relationship matrices constructed either from SNV marker or from the proportion of shared haplotypes across 1cM windows. (see Table 1). The two approaches yield similar estimates, showing that both high-density genotypic data and haplotypic data capture genetic variance.

**Fig. 4.**
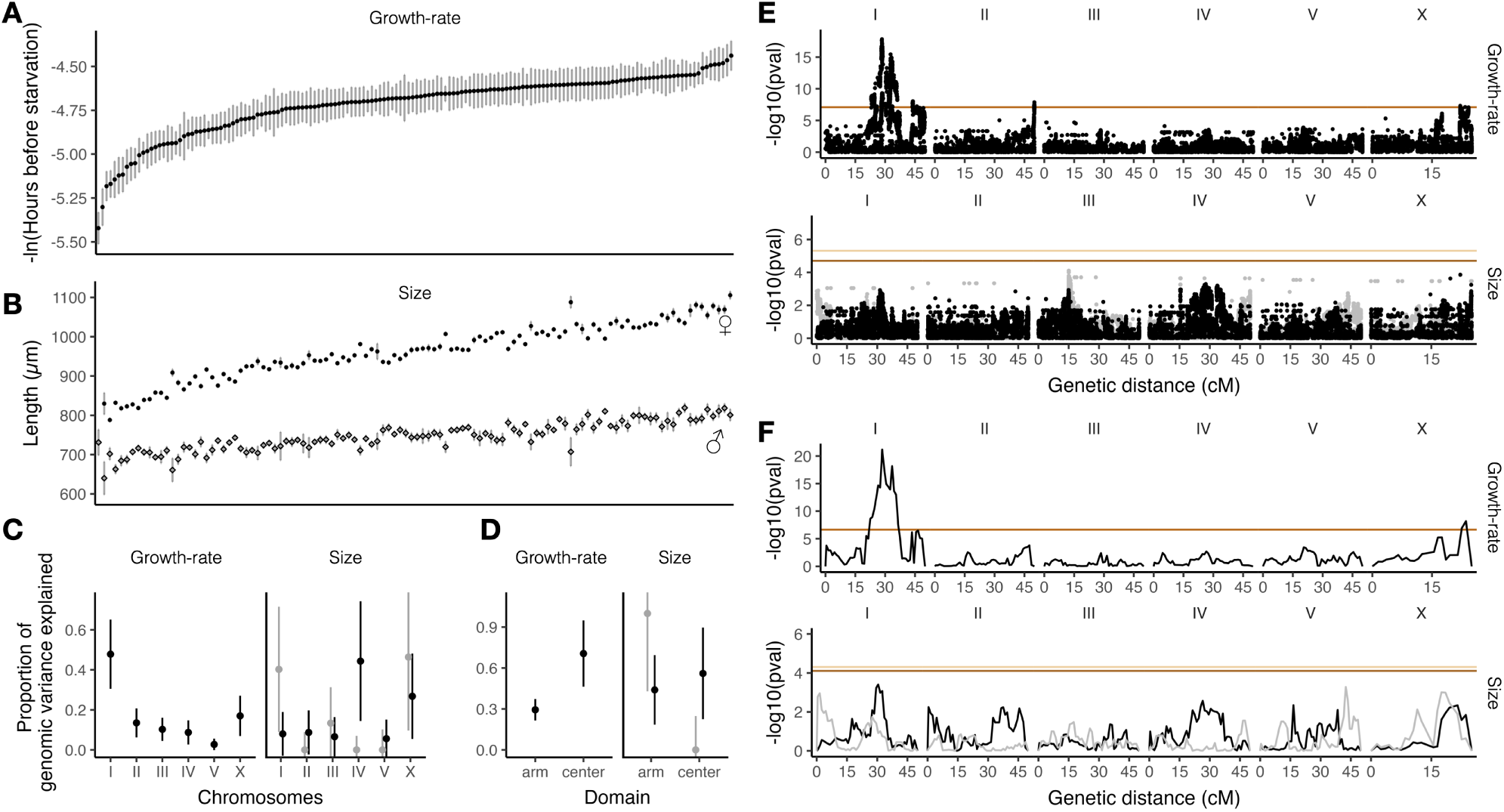
QTL mapping. **A** and **B**. Distribution of trait values for the two phenotypes of interest (mean ± SE). For size, squares indicate male estimates and circles indicate female estimates. **C,D**.Proportion of genomic variance explained by different genomic regions, estimated from a joint model using a multiple GRM. For size, the sexually-convergent component is plotted in grey, and the sexually-divergent component is plotted in black. The variance partitioning was done for the six chromosomes (A) and the recombination domains (B), as defined in Salome Correa et al. (2025); tip domains were pooled with arm domains. **E**. QTL mapping by SNV association test, with the significance threshold from random permutation indicated by the dashed orange line. **F**. QTL mapping by haplotype association test. For size, the sexually-convergent component is plotted in grey and the corresponding significance threshold in light orange. The sexually-divergent component is plotted in black with the corresponding significance threshold in dark orange.

**Table 1.** Genomic heritability for the traits of interest, estimated using a GRM computed from SNV alleles 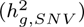 or from shared haplotypes 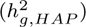. Standard errors are shown in parentheses.

|  | Growth-rate | Sex-conv. Size | Sex-div. Size |
| --- | --- | --- | --- |
| $h_{g,SNV}^2$ | 0.63 (0.05) | 0.25 (0.10) | 0.49 (0.10) |
| $h_{g,HAP}^2$ | 0.67 (0.04) | 0.26 (0.10) | 0.45 (0.10) |

The second trait we investigated is adult size 48 h post-L1, measured for males and females (Figure 4B). We transformed sex-specific body size into sexually convergent and sexually divergent components without loss of phenotypic variance. The sexually convergent axis corresponds to the direction explaining the greatest proportion of variance in the data, whereas the sexually divergent axis is the orthogonal direction (see Methods). Compared to analyzing the sexes separately, this approach explicitly leverages the cross-sex correlation in body size to increase power for detecting genetic variation affecting both sexes similarly (the sexually convergent axis), while also isolating sexually antagonistic genetic variation (the sexually divergent axis). Moreover, the amount of genetic variance along these two orthogonal axes determines the extent to which body size can evolve jointly or independently between the sexes, thereby quantifying the genetic constraints on the evolution of sexual dimorphism. In outcrossing species such as *C. becei*, these two orthogonal axes not only reflect developmental and mutational constraints, but also the action of sex-specific selection shaping standing genetic variation, in contrast to self-fertilizing species where selection on males is limited.

The genetic variances (*σ*^2^) are respectively 0.36 and 0.08 for the sexually convergent and divergent axes, corresponding to a sex correlation of *r_m,f_* = 0.63. This moderately high genetic correlation indicates that the evolution of further sexual dimorphism in size is constrained, but not prevented. Despite lower genetic variance in the sexually divergent component, it shows higher heritability compared to the sexually convergent component (Table 1), because of proportionally lower environmental variance. However, we attribute this pattern to an experimental artifact resulting from environmental covariance between sexes, as both sexes from a given strain are grown together, i.e., size difference between sexes is self-corrected for small variation across Petri dishes.

To test whether genetic variation was uniformly distributed across the genome, we then partitioned the genomic variances across different chromosomes for all traits (Yang et al., 2011). We found that genomic variance is unevenly distributed; for instance, chromosome I explains a large fraction of the genomic variance for multigenerational population growth rate (Figure 4C). We also partitioned genomic variance between chromosomal centers and arms (Figure 4D), as these recombination domains differ strongly in genomic composition, including gene density and nucleotide diversity. Despite centers accounting for a smaller share of the overall genetic diversity, they represent a substantial share of the genomic variance for growth rate and sex-divergent size. This contrasts with *C. elegans*, where genetic variance is biased against chromosomal centers for both fitness-related traits and gene expression (Rockman et al., 2010; Noble et al., 2017). This difference might reflect how selection operates in gene-dense centers under different mating systems: under predominant selfing, recessive deleterious mutations and their linked sites are more effectively exposed to purifying selection, whereas under obligate outcrossing, recessive deleterious variants can remain masked in natural populations but become exposed in the RILs analyzed here.

### QTL mapping

We tested single SNV-marker effects on the two traits of interest using LMMs, accounting for genome-wide relatedness (Figure 4E). At a significance threshold of *α* = 0.05, determined by permutation, we identified two QTLs on chromosome I, one on chromosome II and one on chromosome X, for multi-generational population growth rate. No QTL was detected for size. The absence of detected QTL for size is not surprising as it is well known to be a highly polygenic phenotype.

We also implemented a haplotype-based LMM mapping approach, iterating across genetic positions and testing for haplotypic multi-allelic effects while accounting for relatedness derived from shared haplotypes (Figure 4F). This method detected two growth rate QTLs already identified with the SNV-based approach.

### Correlation of allele’s effect on growth rate and selection during panel derivation

We then investigated whether selection may have occurred during panel derivation. Because our measure of multi-generational growth rate is expected to capture major fitness components, we tested the relationship between allelic effects on growth rate and allele-frequency changes during panel derivation (Figure 5A). For instance, the central QTL on chromosome I is largely composed of SNVs for which the minor allele uniquely tags the founding haplotype “FM.g2”. These alleles are predicted to negatively affect growth rate and, consistently, decreased in frequency in panel *α*, from 25.0% in the founders to 17.7% in the RILs. More generally, the SNV *t*-statistics from the QTL mapping correlate well with allele-frequency changes since the founders (*r* = 0.42, *p* = 0.018, permutation-based). Even when excluding chromosomes harboring QTLs, the correlation remains positive, although only marginally significant (*r* = 0.32, *p* = 0.083, permutation-based). This is notable because several factors are expected to weaken this correlation, including allele-frequency changes caused by drift, the non-linearity between selection strength and *t*-statistics, and the fact that multi-generational growth rate in the RILs likely does not capture all components of fitness, such as male competitive ability.

**Fig. 5.**
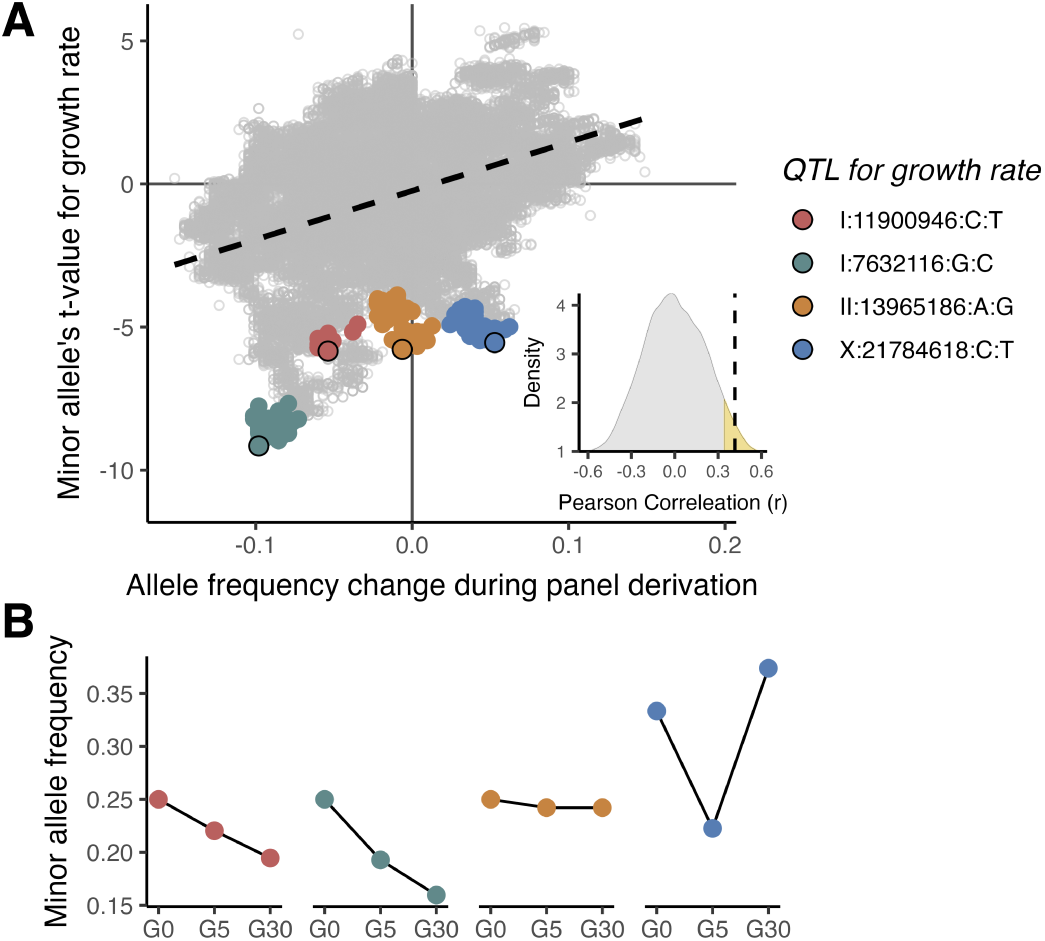
Selection and allelic effects on growth-rate. **A**. Each dot represents the minor allele at a single SNV in the panel *α*. The leading SNV of the four identified growth rate QTL identified in Figure 4E is highlighted by a large colored dot with black borders. Smaller dots highlight SNVs in high linkage disequilibrium with the leading SNVs (*r*^2^ *>* 0.8). Other SNVs are represented in grey. The inset shows the null distribution of correlation coefficients obtained by 10^4^ permutations (single-sided p-value = 0.018). For each permutation, random allele frequency changes were obtained by permuting the founding haplotype frequencies in the RILs within each chromosome (i.e., relabeling the colors in Figure 2B) and extracting the allele frequencies. This method conserves the allele frequency variance and the linkage structure in the permuted datasets. The relationship is presented separately for each chromosome in Figure S8. **B**. Minor allele frequencies of each QTL in the founders (G0), in the population after the outcrossing phase (G5), and in the RILs following inbreeding (G30). Estimates were obtained by averaging the allele frequencies at the hundreds of linked SNVs composing the QTL (i.e., the colored dots in panel A).

The direction of selection matches the sign of the *t*-value for all QTLs except the one on the X chromosome. This QTL is also the only one that shows a striking reversal in its allele-frequency trajectory between the outcrossing and inbreeding phases of panel derivation (Figure 5B). For instance, a trade-off between male competitive ability and female fertility could underlie such a reversal, as male-male competition is absent during the inbreeding phase, which involves random mating pairs. Future studies quantifying sex-specific competitive fitness could improve our understanding of how the genetic architecture of fitness shapes selection in outbred nematodes.

## Discussion

The *C. becei* RILs (beRILs) presented here represent the first large-scale RIL panel –indeed, the first genetic reference panel – in a gonochoristic *Caenorhabditis* species. We demonstrate their suitability for quantitative genetics and QTL mapping, highlighting the potential of these panels for researchers interested in comparative genetics and in traits that are particularly relevant to outcrossing species. The beRILs are compared to a selection of other broad-application *Caenorhabditis* RIL panels in Table 2. The beRILs are outstanding in almost every category, with the important exception of LD, where CeMEE remains unmatched.

**Table 2.**
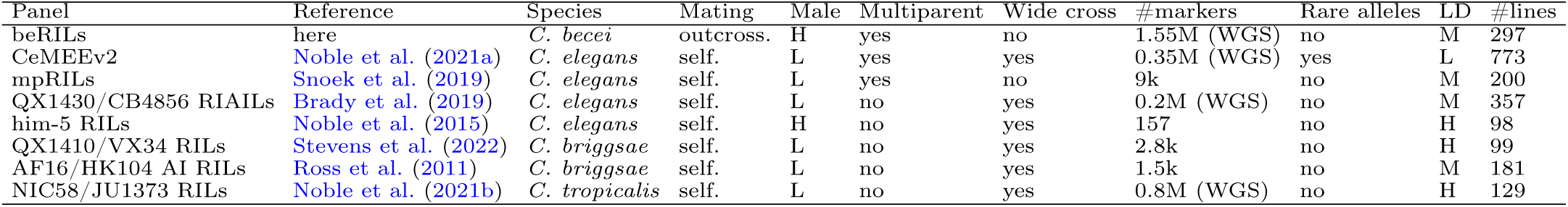
Comparison of recombinant inbred line panels across *Caenorhabditis* species. ”Male” indicates whether the male frequencies within a line are high. “Wide cross” indicates whether the population founders originate from very distant locations or from a single location. “#markers” is the number of available informative markers reported after quality control. It depends on the diversity segregating in the panel and whether targeted genotyping or whole-genome resequencing (WGS) was used. “Rare alleles” indicates whether the minor allele distribution is skewed toward low frequencies (*<* 10%). “LD” indicates whether the linkage disequilibrium is high. “#lines” is the number of derived lines that are genotyped. Abbreviation: H = high, L = Low, M = moderate.

The beRILs panels sample six haploid genomes from a natural population found in the tropical forest of Barro Colorado Island (BCI), Panamá (Sloat et al., 2022). Only a few generations and one round of cryopreservation separate the sampling of the natural isolates from the crosses that initiated the panels. Thus, the components of the genetic architecture observed in these RIL panels are likely to be relevant to the local population from which they originate. This contrasts with most *Caenorhabditis* RIL panels, which originate from intercrosses of divergent and geographically distant strains and are therefore prone to genetic incompatibilities that may not reflect the structure of many natural populations (Colomé-Tatché and Johannes, 2016). The closest analog may be a multiparental panel in *C. elegans* whose four haplotypes derive from two localities in France separated by a hundred kilometers (Snoek et al., 2019). Phenotypic analyses of panel *α* show that a single fig harbors a lot of standing variation for important traits. Some will reflect recessive alleles, that are often masked in nature, but not completely, as illustrated by the presence of large regions of ROH in some founders. The population in a fig represents diversity introduced to that habitat patch by a small number of individuals (Richaud et al., 2018; Sloat et al., 2022; Rockman et al., 2026). Despite this presumed history of bottlenecking and biparental inbreeding, the panel harbors genetic variation on every chromosome that affects multigenerational growth rate in the lab, including several significant QTLs whose effects are mirrored by slight shifts in allele frequencies during the construction of the RIL panel. The RILs *α* also harbor heritable variation for body size, with female length differing among strains by hundreds of microns, the longest strains on average more than 30% longer than the shortest. Though much of the variation in body size is shared by males and females, there is a significant and heritable component of sexually divergent body size variation, the fuel for evolutionary change in sexual dimorphism (Cox and Calsbeek, 2009; Kaufmann et al., 2023).

Though haplotype frequencies changed during strain construction in a manner consistent with directional selection on the population-growth-rate QTLs, the overall magnitude of haplotype frequency changes during panel derivation was modest, and no founding haplotype was lost. Here again, we see important differences with panels developed for selfing species, where strong allele frequency distortions are common. These distortions are due primarily to the segregation of Medea and peel alleles that have caused gene drive effects in the construction of RIL panels in *C. elegans* (Seidel et al., 2008), *C. briggsae* (Ross et al., 2011; Ben-David et al., 2021), and *C. tropicalis* (Noble et al., 2021b). This phenomenon is expected to be a common feature of crosses between divergent selfing isolates (Rockman, 2025), and, although Medea alleles also occur in *C. becei* (Salome Correa et al., 2025), they appear not to segregate among the founding haplotypes of our panel. *C. elegans* panels also show allele frequency distortions due to sweeps by lab-adaptive alleles (Noble et al., 2021a) and likely to outbreeding depression, negative epistatic interactions that are expected to characterize distant crosses among selfers, whose genomes have coadapted loci thanks to their rare outcrossing and extended linkage disequilibrium (Dolgin et al., 2007; Snoek et al., 2014; Gimond et al., 2013); our panel escapes these effects. Though inbreeding depression has been the scourge of gonochoristic *Caenorhabditis* genetics, our genotype data imply the absence of recessive lethals in our founding haplotypes and no excess of heterozygosity after 25 generations of sib-mating.

Gonochoristic species are far more diverse than their predominantly selfing relatives (Jullien et al., 2019), a pattern that reaches extremes in *Caenorhabditis* (Dey et al., 2013; Teterina et al., 2023). The beRILs, sampling the genomes of three worms from a single location, capture at least 2.2 million genetic variants, roughly the same number that segregates in the entirety of the global *C. elegans* population, as represented by more than 600 genomes from every continent but Antarctica (Lee et al., 2021). The *C. becei* panel harbors about five times the number of informative markers reported in the CeMEE, the most diverse and recombinant RIL panel in *C. elegans* (Noble et al., 2021a). Moreover, most variants in the beRILs segregate at intermediate frequencies, which increases the power to detect QTLs.

Although the founding haplotypes were allowed to recombine for only a limited number of generations, appreciable mapping resolution was achieved; causal variants are expected to lie within less than 1 cM of the peak. However, fine-mapping causal variants within the low-recombining chromosomal centers remains challenging. This difficulty is well known in *Caenorhabditis* (e.g., Alkan et al. (2024)), and reduced resolution in chromosomal centers is still observed in the CeMEE even after more than 140 generations of recombination (Noble et al., 2021a). At the same time, the genetic map of the panel suggests an absence of large segregating inversions or other recombination-suppressing features.

An advantage of having clearly identifiable haplotype blocks is the straightforward implementation of haplotype-based methods for quantitative genetics and mapping. This framework will also facilitate the future incorporation of structural variants into the dataset with minimal resources: sequencing a handful of RILs with long reads and assigning structural variants to their founding haplotypes will be sufficient to impute them across all RILs.

Another major promise of the beRILs is the possibility of coupling quantitative genetics with RILs and experimental evolution with the outcrossing populations. In *C. elegans*, this approach has already been used, for example, to compare genetic variance 693 for fitness and patterns of allele frequency change across genomic intervals (Parée et al., 2025), or to predict the direction of phenotypic evolution (Mallard et al., 2023).

In contrast, many experimental evolution studies lack the resources to link individual genotypes to phenotypes and are therefore restricted to population-level observations (Kofler and Schlötterer, 2014).

Together, these results establish the beRILs as a powerful resource for dissecting the genetic architecture of complex traits in an obligately outcrossing *Caenorhabditis* species. Beyond their immediate use for QTL mapping, these panels lay the groundwork for comparative analyses across mating systems and for investigating the evolutionary forces maintaining genetic diversity in the genus *Caenorhabditis*. Finally, these panels represent a new superlative for experimental quantitative genetics in animals: a panel with millions of SNVs in characterized haplotypes with intermediate frequencies, similar to resources available for *Drosophila*, but here in a trivially cryopreservable animal with a two-day generation time.

## Acknowledgments

This work was supported by grants R01GM121828, R01HG013015, and R35GM141906 (MVR) from the National Institutes of Health, and New York University Dean’s Undergraduate Research Fund awards (SV). We thank NYU GenCore, supported by the Zegar Family Foundation, for sequencing support, and NYU HPC for computational support. We thank Sol Sloat for help in the lab and Luke Noble for discussion.

## Supplementary Information

### Supplementary Figures

**Fig. S1.**
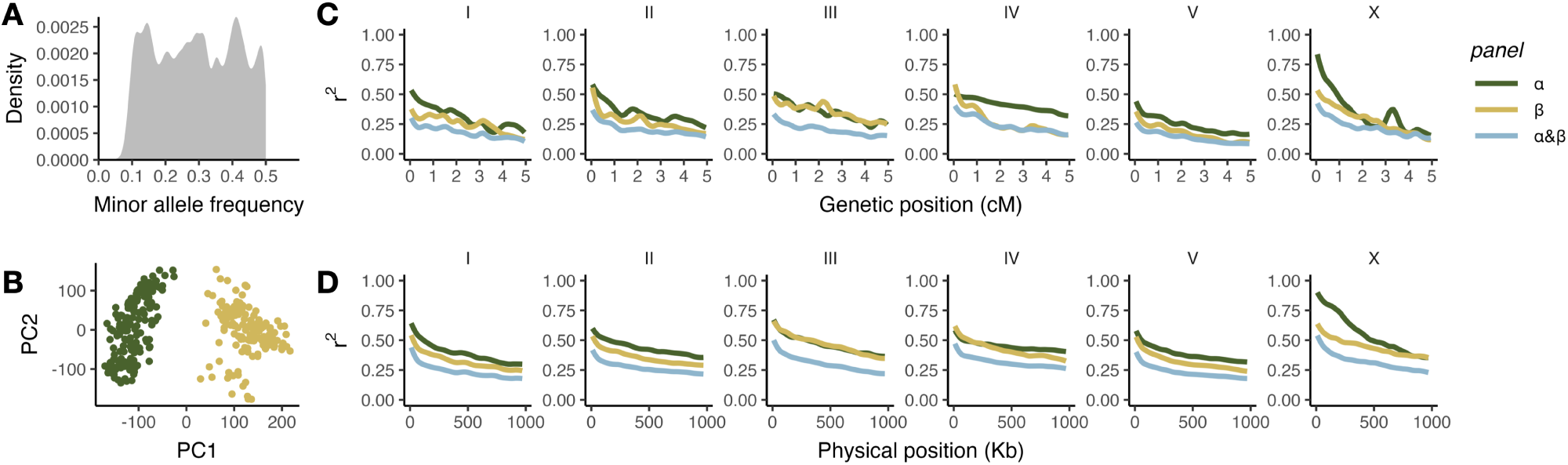
Population structure and linkage disequilibrium decay by chromosome. **A.** Minor allele frequency spectrum **B.** Principal component analysis (PCA) of the RILs genotypes. **C.** Linkage disequilibrium (LD, *r*^2^) as a function of genetic distance between markers. **D.** Linkage disequilibrium (LD, *r*^2^) as a function of physical distance between markers.

**Fig. S2.**
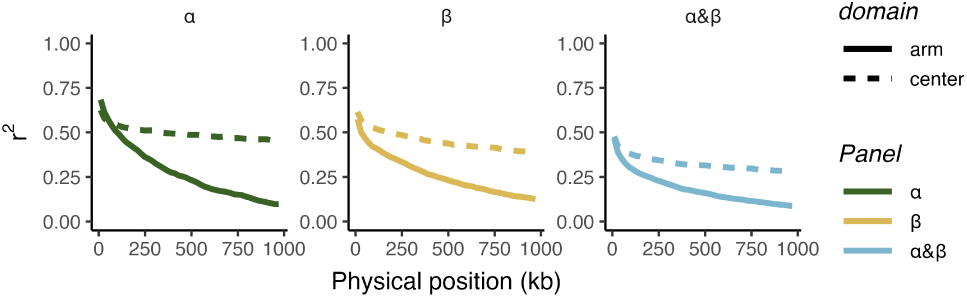
Autosomal linkage disequilibrium decay by recombination domains.

**Fig. S3.**
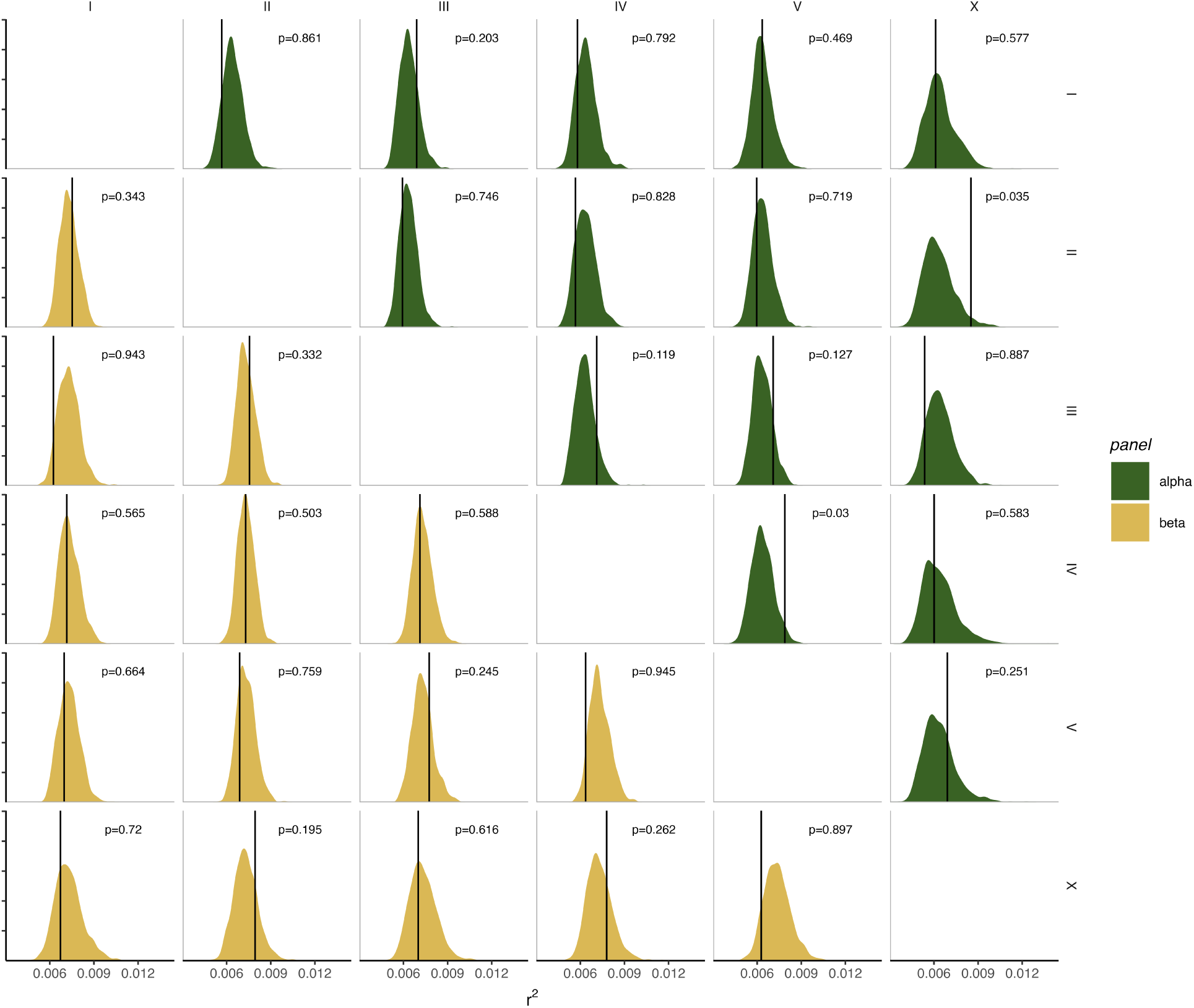
Inter-chromosomal linkage. Linkage disequilibrium (*r*^2^) across chromosome pairs was calculated. To reduce the number of pairs of variants and the computational burden, dataset pruned for intra-chromosomal linkage *r*^2^ *>* 0.8. For each possible chromosome pair within panel, a null distribution of mean inter-chromosmal linkage was obtained by randomly permuting 1,000 times the chromosomes across RILs. The vertical black line show the observed values. None of the chromosome pairs shows greater linkage than expected (one-sided p-value; Bonferroni threshold = 0.05*/*30 = 1.6 × 10*^−^*^3^).

**Fig. S4.**
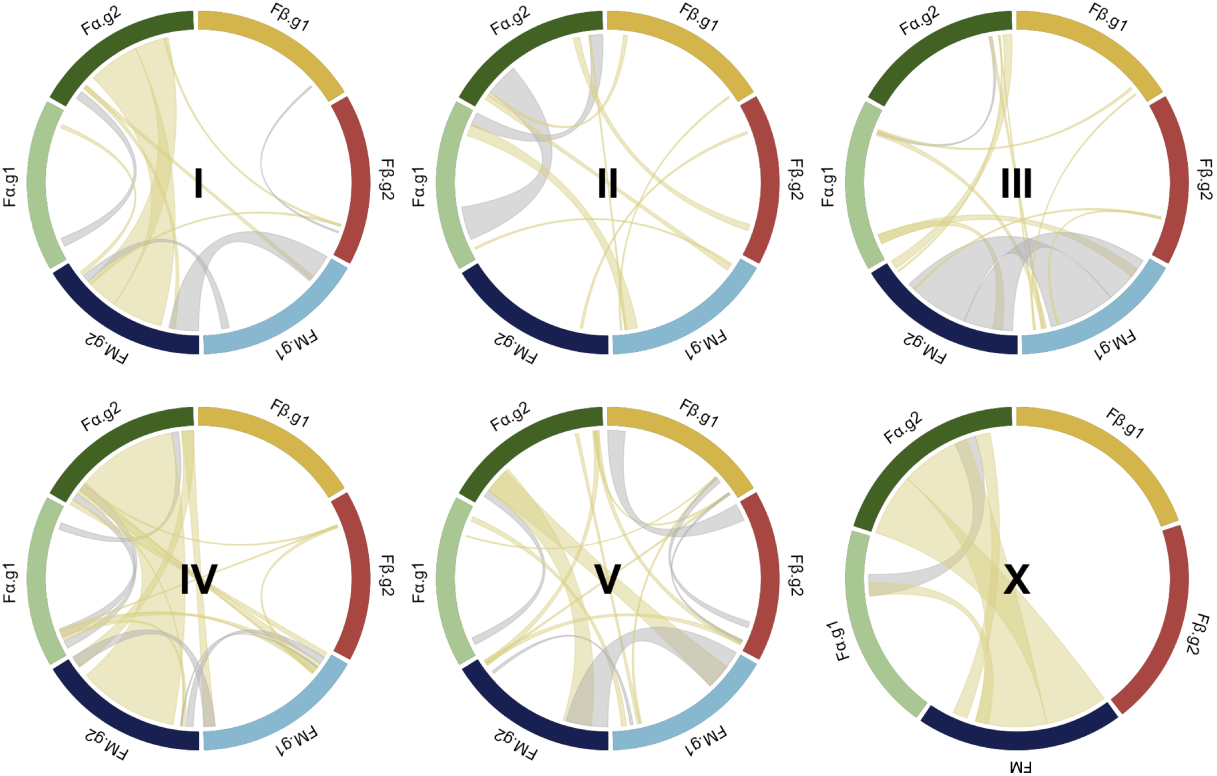
Homology between founder haplotypes. For each chromosome, the chord plot segments represent the three pairs of founding haploid genomes, with colors indicating their founder of origin (green: *Fα*, orange: *Fβ*, blue: *FM*). Haplotype blocks larger than 2cM that are identical between two founding haplotypes are highlighted by lines: beige when shared between distinct founders and grey when shared within a founder (i.e., homozygosity).

**Fig. S5.**
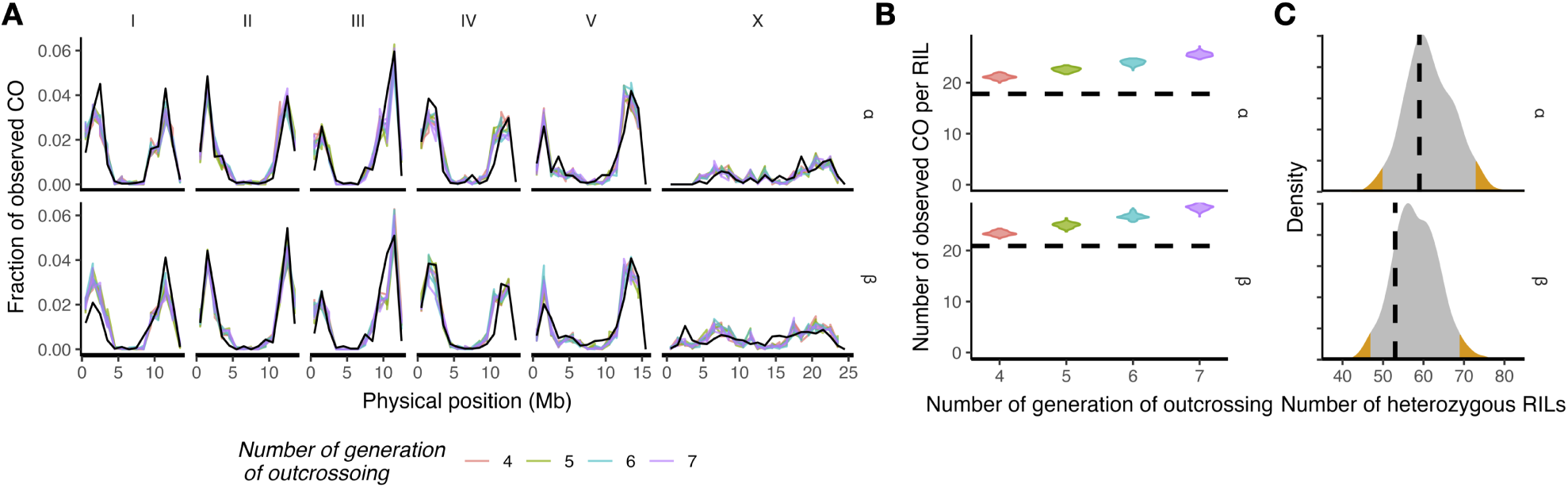
Neutral simulations. Distribution of recombination breakpoints (**A**) and average number per RIL (**B**), as well as the number of RILs with heterozygous genotypes (**C**) in the simulated datasets. In the simulations, RILs were derived from populations undergoing varying numbers of outcrossing generations (*g*outcross = 4–7) and population sizes (*N* = 400 or 4000). Simulations with different population sizes are pooled in (A) and (B), and all parameter combinations are pooled in (C). Experimental observations are indicated by black lines in each plot.

**Fig. S6.**
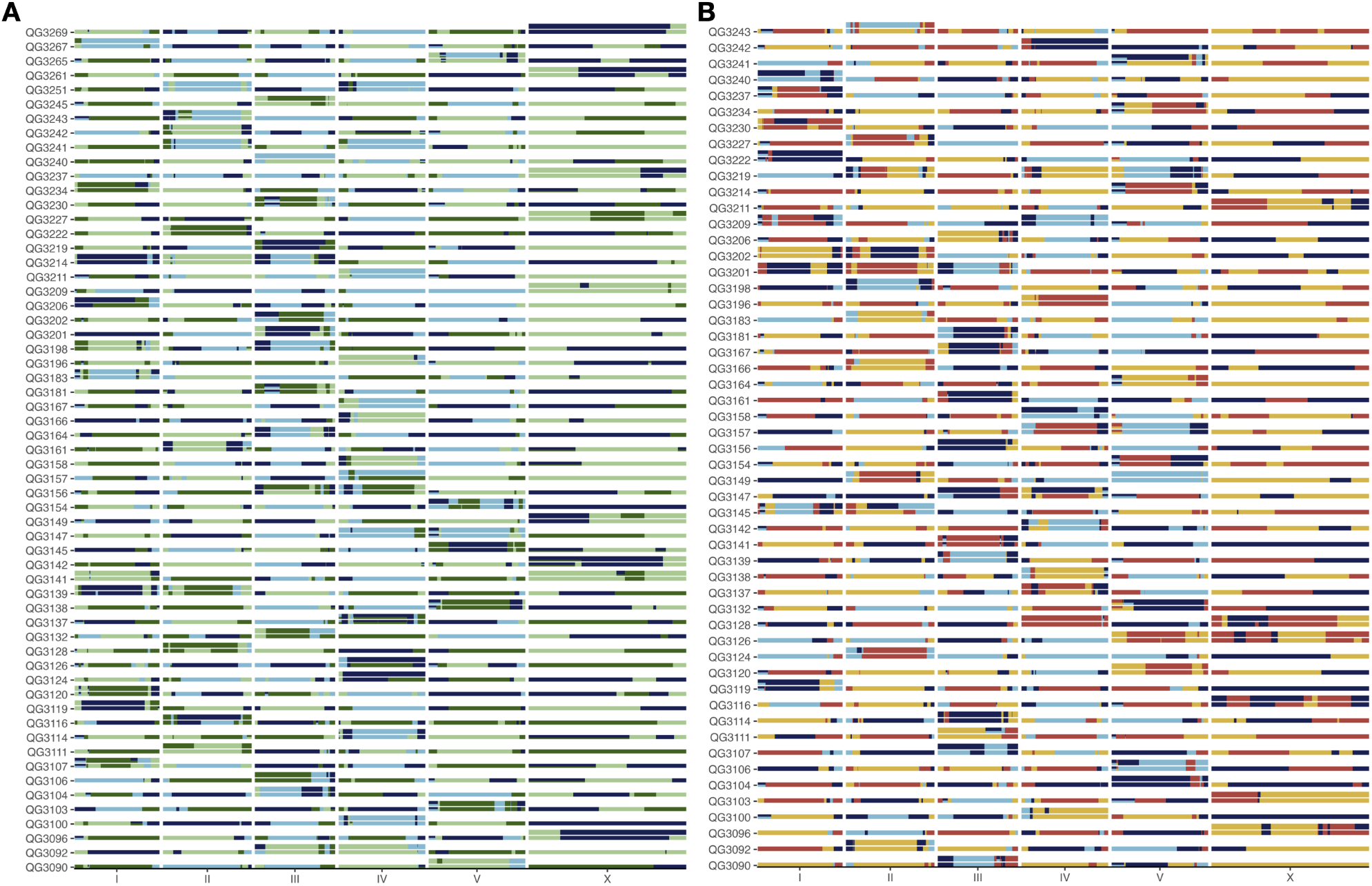
Diplotype of heterozygous RILs. For each RIL bearing a heterozygous diplotype block, the haplotype blocks are represented. When a chromosome is completely homozygous, only a single haploid copy is shown. When a chromosome contains heterozygous segments, the two reconstructed divergent haploid copies are shown. The color scheme is the same as in Figure 2.

**Fig. S7.**
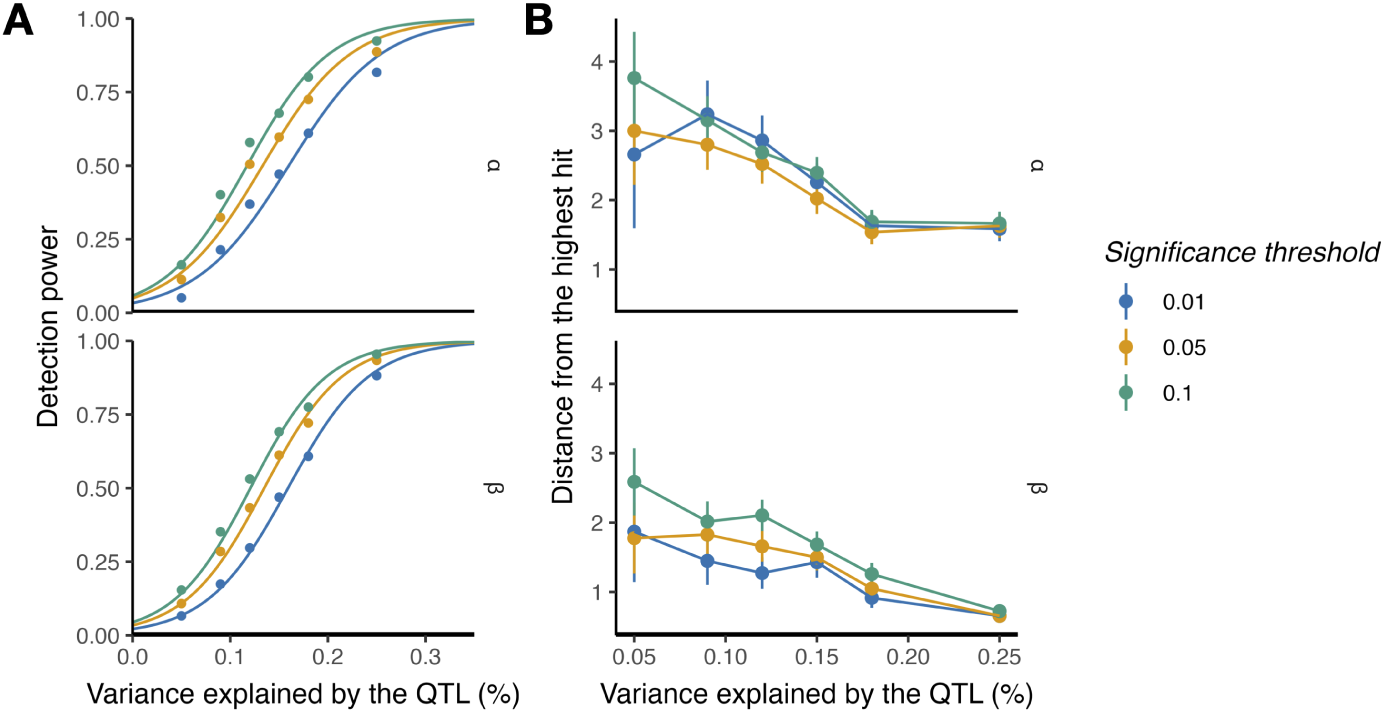
Power analysis using single panel. See legend of Figure 3.

**Fig. S8.**
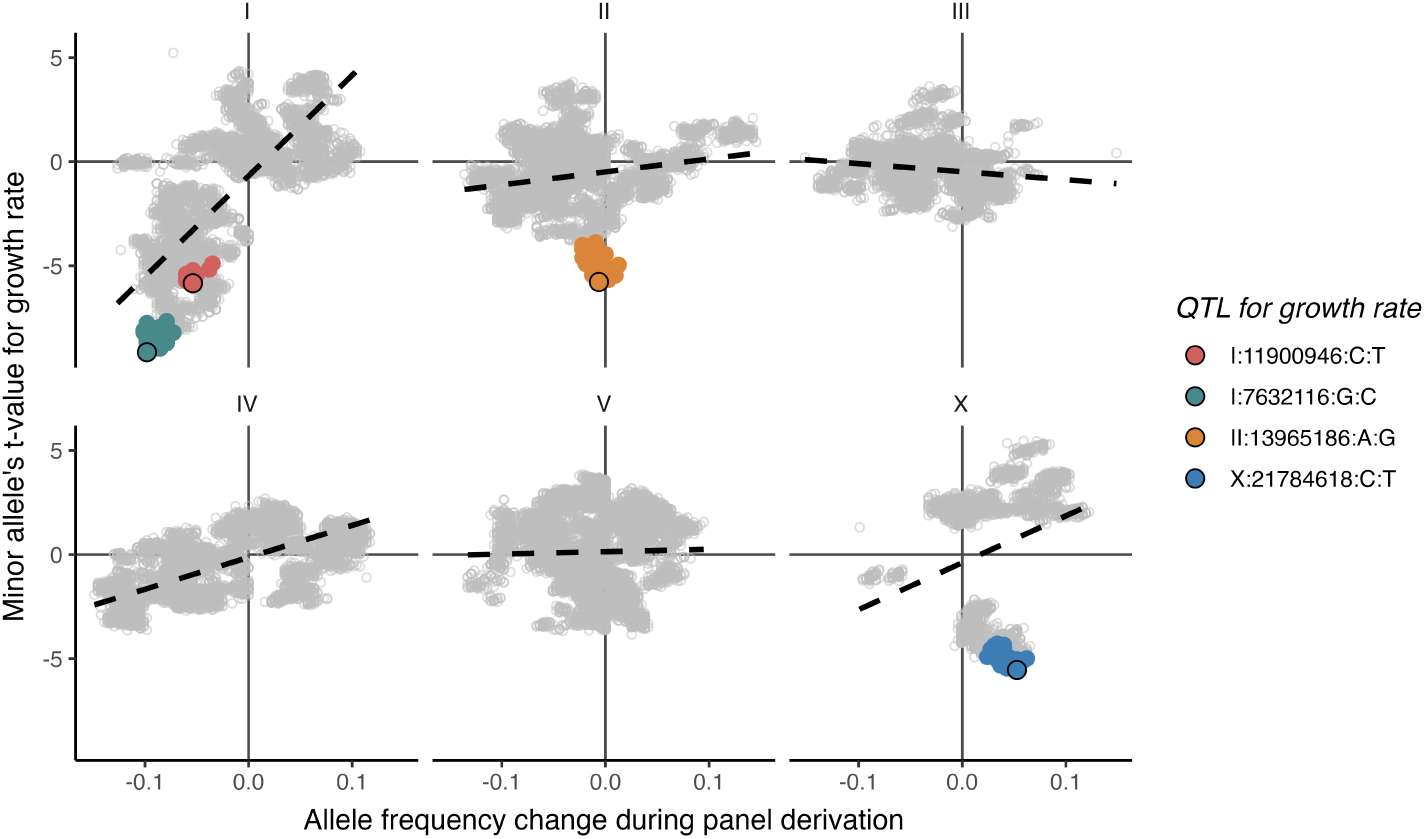
Selection and allelic effects across chromosomes.

### Supplementary Tables

**Table S1.**
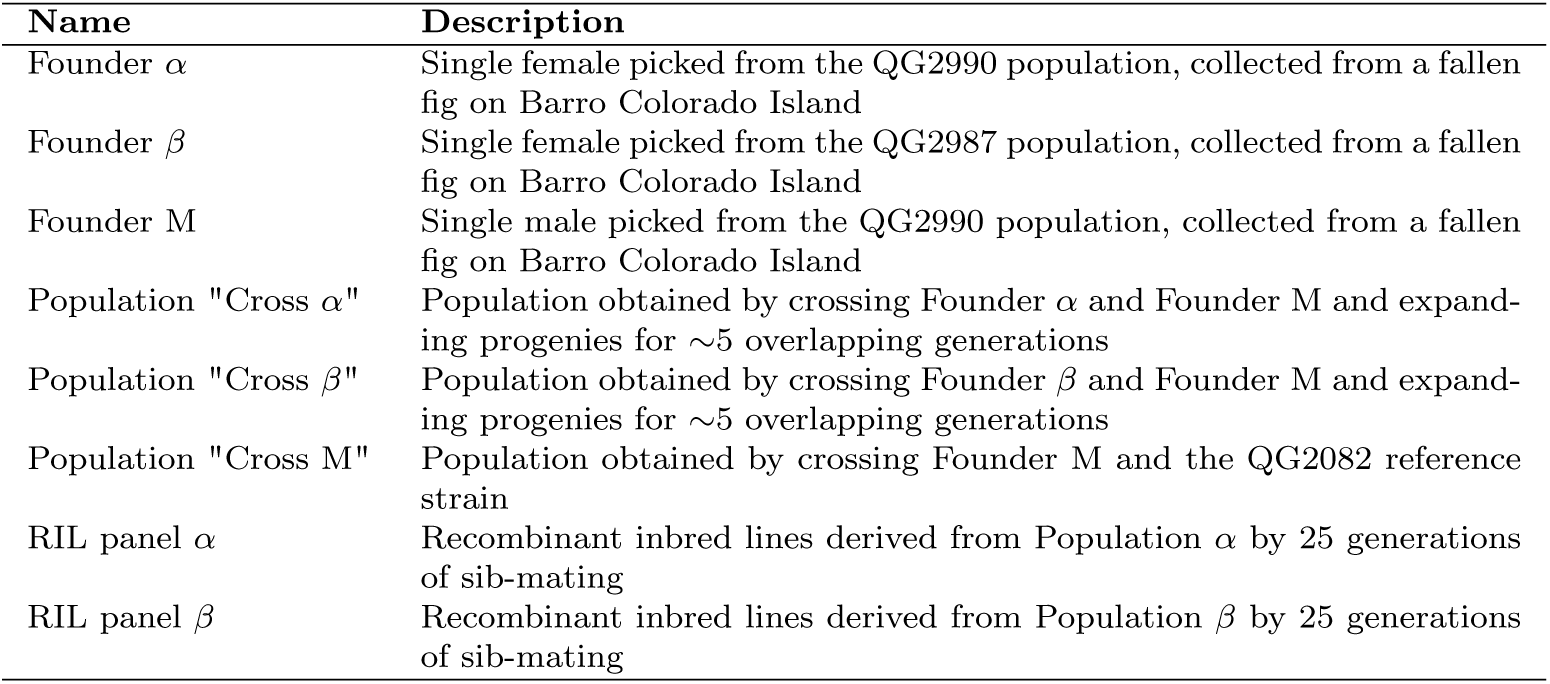

**Table S2.** *f_ROH_* is the inbreeding coefficient measured as the proportion of 1 cM non-overlapping genomic intervals without heterozygous SNV. For the male founder specifically (FM), the X chromosome is excluded from this analysis as it is hemizygous.

| Founder | $f_{ROH}$ | Homozygous variants (%) |
| --- | --- | --- |
| Fα | 0.22 | 62.6 |
| Fβ | 0.08 | 50.5 |
| FM | 0.30 | 68.1 |

## Appendix: haplotype reconstruction

To reconstruct founder haplotypes, call them in the RIL samples, and filter variants, we developed a custom method tailored to our specific experimental design. Scripts and relevant datasets are available at the associated Github repository. First, the likelihood of observing allelic count in pool-sequenced populations expanded from crosses was calculated for every possible founder’s diploid genotypes. The second step involves inferring the six founding haploid genomes using the diploid genotype likelihood, linkage information present in RILs, and the panel design. To infer the ‘backbone’ of founding haplotypes, the stringent dataset containing only the most confident SNVs was used (see methods). The third step is to call founder haplotype blocks across the RILs. The fourth step uses the inferred RIL haplotypes to identify valid variants. The fifth step is to call the haplotypes a second time, but now using all the variants.

## 1. Founder genotype likelihood probability

The initial crosses that founded the two RIL panels were pool-sequenced prior to cryopreservation (cross *α* and *β*). The cross between the male founder (FM) and the QG2082 reference strain was also pool-sequenced (cross *M*). For every possible combination of diploid genotypes of the three founders, the expected true allele frequency in the crosses, denoted *θ*, can be derived. For example, under the hypothesis that founders *α*, *β*, and *M* have genotypes 0*/*0, 0*/*1, and 1*/*1, the expected allele frequencies *θ* are 0.5, 0.75, and 0.5 in crosses *α*, *β*, and *M*, respectively. Given the allelic counts, we can compute the log-likelihood of the founder genotype under each scenario of founder genotype combinations. Let *k* denote the number of observed alternative allele reads and *n* the total read depth. The underlying true allele frequency is denoted *θ*, and the sequencing error rate *ε* is set to 10*^−^*^4^. The probability of observing an alternative allele is:

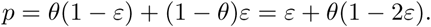

And, the observed count follows a binomial distribution conditional on *p*:

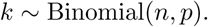

The log-likelihood of *θ* given the data is therefore:

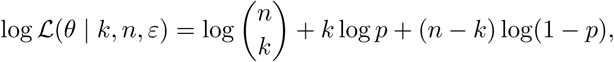

The method was implemented in R (see FounderGenotype_logLikelihood.R).

## 2. Phasing the founding haploid genome

The haplotype blocks segregating in the RILs were used to help reconstruct and phase the six founding haploid genomes while maximizing the log likelihood of founders’ diploid genotypes calculated above. A limitation of this approach is that if a founding haplotype was lost before RIL derivation, it cannot be inferred. Such a possibility would be highlighted by a genomic interval with lower log likelihood values, larger allele frequency deviation between crosses and RILs, and recombination breakpoints wrongly inferred, leading to distortion relative to the genetic maps. We did not detect such regions.

The reconstruction of the founding haploid genomes, was implemented using a custom algorithm in R (see 00_beceiFounders_phasing.R). We used an overlapping-window-based approach, identifying segregating haplotype blocks and assigning them to the different founders based on the constraint of the experimental design and the founders’ diploid genotype inferred from the pool-sequenced populations. For each 500-SNVs sliding window and each RIL panel (*α* or *β*), RILs sharing the same exact haplotype block were grouped. The four founder haplotype blocks segregating in each panel were assumed to be the four most common haplotypes, as recombination is expected to be rare within these small windows (typically <1 cM). Since haplotype similarity between founders may reduce the number of segregating founder haplo-types, only haplotypes present in at least 5 RILs were considered plausible founder haplotypes. This threshold confidently excludes any rare haplotype arising from recombination or genotyping error. Each founder was then matched to a pair of haplotype blocks that maximizes the sum across SNVs and founders of the founder’s diploid genotype log-likelihood (see previous section). Founder M was only matched to haplotypes shared between both panels, as expected from the cross scheme. At this stage, some founder haplotype blocks might be misinferred. To verify consistency, we assessed whether overlapping windows inferred the consistent haplotype pairs for all founders. When two overlapping windows were incompatible, the inference was repeated after merging the incompatible windows to provide additional information for haplotype block assignment. This step successfully resolved all detected incompatibilities.

Once founder haplotype blocks were inferred within each window, the windows needed to be phased relative to one another. For each pair of adjacent windows, we tested both possible phasing orientations and selected the one that minimized the number of recombinant events among RILs.

## 3. Calling haplotype blocks in RILs from the stringent dataset

Founder haplotype blocks were then called in each chromosome for a given high-quality RIL, by adapting a custom R script from Parée et al. (2025). Briefly, many windows of varying sizes are created across the genome. For a given genomic window, the founding haplotype or the combination of founding haplotypes (in case of a heterozygous region), minimizing the Hamming distance from the observed RIL genotypes is found.

The threshold to validate a match is 99% for a homozygous region, and 97.5% for a heterozygous region, because heterozygous alleles are not always observed accordingly due to sampling, particularly at low depth. Then, the boundaries of each window are extended until the hamming distance between RIL and founder haplotypes starts to increase, indicating a recombination breakpoint. The custom algorithm can be found at 01_haploCall_RILs.R.

## 4. Incorporating variants that are coherent with the inferred haplotype blocks

The inferred RILs haplotype blocks were then used to search for any valid genotype that have been filtered out from the high-quality dataset. A custom R script was used to iterate across all SNVs (02_filterHaplotypeConsistentVariants.R). For a given SNV, the genotypes are grouped based on the founding haploid genome identity at this genomic position. If a unique allele was found for each founding haploid genome group, it means that the SNV is consistent with the haplotype blocks already inferred in the RIL, and it was kept. Alternatively, if more than one allele was seen in at least one founding haploid genome group, the SNV was filtered out as it would imply multiple versions of the same haplotype. We allowed for 2 genotyping errors across all RILs.

## 5. Calling haplotype blocks with all SNVs and imputing missing SNVs

The haplotype calling was repeated (section 3), but using all the validated SNVs in the previous step (03_haploCall_RILs2.R). The inferred haplotypes were then used to impute missing SNVs using inferred haplotypes (04_inpute_RILs_missing_SNPs.R).

